# Anticipating Protein Evolution with Successor Sequence Predictor

**DOI:** 10.1101/2024.06.08.598054

**Authors:** Rayyan Tariq Khan, Pavel Kohout, Milos Musil, Monika Rosinska, Jiri Damborsky, Stanislav Mazurenko, David Bednar

## Abstract

The quest to predict and understand protein evolution has been hindered by limitations on both the theoretical and the experimental fronts. Most existing theoretical models of evolution are descriptive, rather than predictive, leaving the final modifications in the hands of researchers. Existing experimental techniques to help probe the evolutionary sequence space of proteins, such as directed evolution, are resource-intensive and require specialised skills. We present the Successor Sequence Predictor (SSP) as an innovative solution. Successor Sequence Predictor is an *in silico* method that mimics laboratory-based protein evolution by reconstructing a protein’s evolutionary history and suggesting future amino acid substitutions based on trends observed in that history through carefully selected physicochemical descriptors. This approach enhances specialised proteins by predicting mutations that improve desired properties, such as thermostability, activity, and solubility. Successor Sequence Predictor can thus be used as a general protein engineering tool to develop practically useful proteins. The code of the Successor Sequence Predictor is provided, and the design of mutations will be also possible via an easy-to-use web server https://loschmidt.chemi.muni.cz/fireprotasr/.

**Graphical Abstract:** 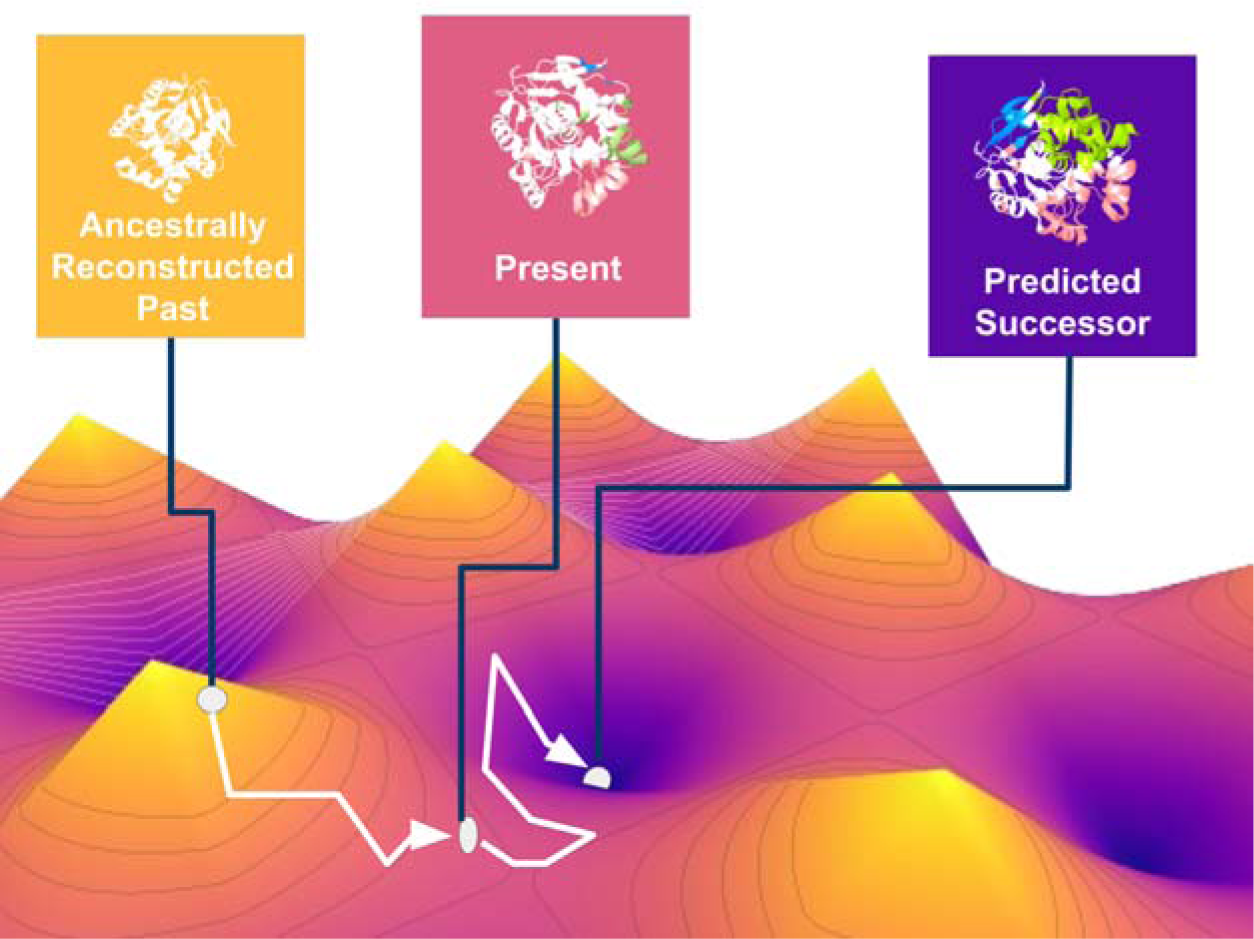

Tracing Evolution’s Pathway to the Future

## Introduction

Evolution is a general term that describes the changes in inherited traits of biological entities through successive generations, generally in response to environmental changes (Hall & Hallgrimsson, 2014). While it can be modelled or described at many levels of biological organisation and varying levels of accuracy, for this study, we will focus on protein evolution.

Protein evolution can be reduced to two key steps: amino acid mutation and the fixation of the mutated protein in a population (Gillespie, 1994; Kimura, 1985). An individual mutation may result from errors in DNA replication during cell division, exposure to mutagens, or a viral infection. The probability of fixation of this new mutation in the population depends on the fitness effect of the mutation itself. The new variant can be neutral, deleterious, or beneficial. While this two-step model is useful, it is only descriptive and not predictive (Nosil et al., 2020). For this reason, it cannot be used to predict upcoming mutations in the future and their fixation probability. Directed evolution techniques are generally used to engineer a protein and possibly understand the effect of various mutations on a protein and their fixation probabilities.

Directed evolution techniques allow a user to probe a protein’s evolutionary space. They are used to improve protein characteristics and, sometimes, even to confer new characteristics onto a protein (Arnold, 2019) by selecting or screening many variants. The markers for improvement in protein characteristics due to induced mutations can be taken as a proxy for fixation probabilities of the induced mutation in a natural environment if it occurs without human intervention. While this model has not been framed in such a way previously, it closely models the concepts of classic Darwinian/positive selection (Fay & Wu, 2000).

However, directed evolution experimental techniques require specialised skills and are both time and resource-intensive. Thus, any in silico technique for predicting and mimicking laboratory-based protein evolution would be of great use for the design of proteins with novel properties. As of this writing, we have only come across one technique, Proseeker, which uses physicochemical characteristics and structure to pick sequences that have higher probabilities of evolving a desired function (Raven et al., 2022). However, the technique was designed specifically for binding proteins. It uses smaller peptide sequences (13 amino acids), and it does not filter AAindices, i.e., physiochemical descriptors (Kawashima, 2000), rather it uses all available AAindices. This leaves room for refinement by selection of more useful indices.

Here, we propose a novel method called Successor Sequence Predictor (SSP), which can mimic laboratory-based protein evolution. It reconstructs the evolutionary history of a protein sequence and then suggests amino acid substitutions based on trends observed in the evolutionary history of the protein when projected through the lens of various, carefully selected, physicochemical descriptors. Introducing the predicted mutations would enhance specific protein properties. For example, if SSP is used on a protein that in the history of its evolution was experiencing a selection pressure towards becoming more thermostable, the predicted substitutions will most likely make the mutant protein even more thermostable, and likewise for other physicochemical properties of the protein. We describe the method in detail and then conduct its critical validation against five different experimental data sets targeting properties such as thermostability, activity, and solubility. A dataset of amino acid sites that were determined to be positively selected by various evolutionary sequence analysis methodologies was also incorporated in the validation (Slodkowicz & Goldman, 2020).

## Materials and Methods

### Selection of AAindices

Nine AAindices were manually selected after consideration, to reflect a variety of possibly relevant physiochemical descriptors (**Table 1**). While the AAindex stores many more indices, they were considered inappropriate due to factors such as redundancy or context-specific physiochemical descriptions. Correlation analysis ensured that the nine selected indices had significant differences (**Figure 1**), and while molecular weight and residue volume indices were similar, they were retained due to the slight nuances of how they evaluated different amino acids.

**Figure 1.**
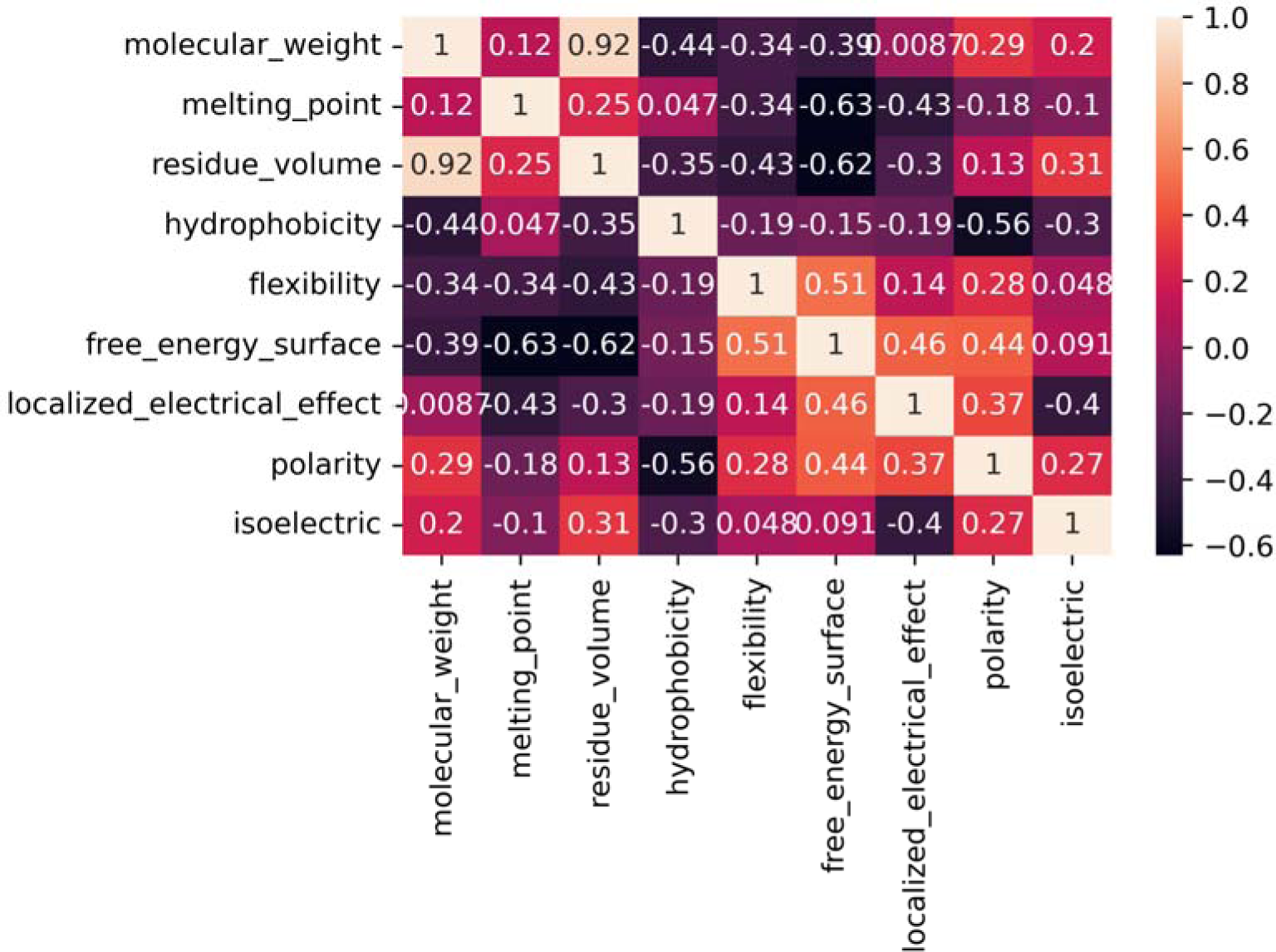
Pearson correlation matrix of selected AAindices. The correlation coefficients are colour-coded from dark purple at −0.7 to off-white at 1.0. The indices are summarised in **Table 1**.

**Table 1.** The AAindices used to analyse amino acid evolution. The correlations among the individual indices are presented in **Figure 1**.

| Index | Property | Reference |
| --- | --- | --- |
| FASG760101 | Molecular weight | Fasman, 1989 |
| FASG760102 | Melting point | Fasman, 1989 |
| GOLD730102 | Residue volume | Goldsack-Chalifoux, 1973 |
| WOLR790101 | Hydrophobicity index | Wolfenden et al., 1979 |
| BHAR880101 | Average flexibility indices | Bhaskaran-Ponnuswamy, 1988 |
| BULH740101 | Transfer free energy to the surface | Bull-Breese, 1974 |
| FAUJ880108 | Localised electrical effect | Fauchere et al., 1988 |
| ZIMJ680103 | Polarity | Zimmerman et al., 1968 |
| ZIMJ680104 | Isoelectric point | Zimmerman et al., 1968 |

### Successor Sequence Predictor Workflow

The workflow of the Successor Sequence Predictor is summarised in Figure 2. For a given target protein, its FASTA sequence is used to mine a dataset of homologous sequences using BLAST (Johnson et al., 2008) while maintaining sequences with identity towards the target sequence between 30-90%. The length filter maintains sequences with 80-120% of the target protein’s length. The remaining sequences are clustered using USEARCH with 90% sequence identity, and a single sequence is randomly selected from each cluster.

**Figure 2.**
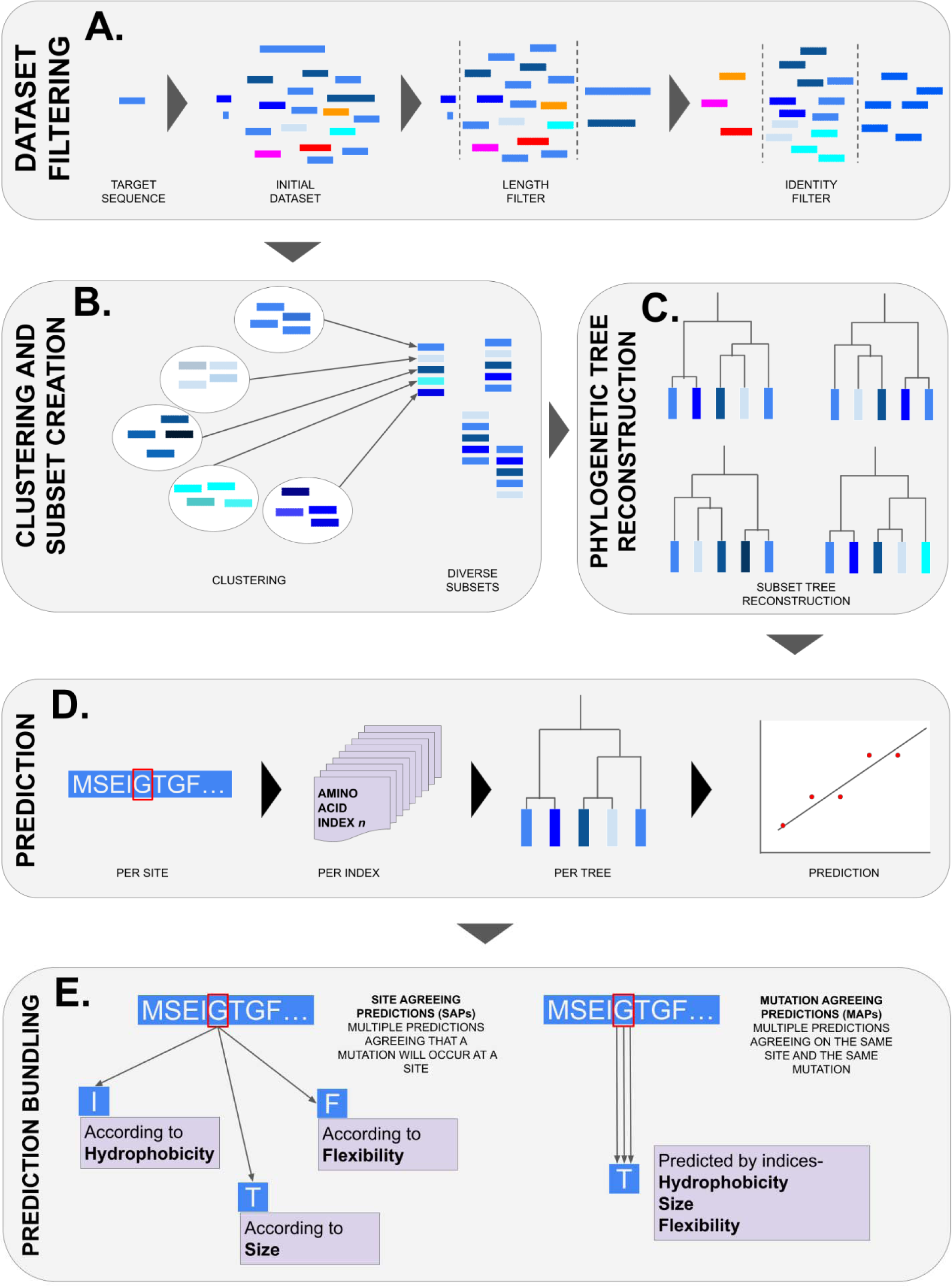
A generalised overview of the Successor Sequence Predictor (SSP). (**A**) Initial curation and filtering of the target protein’s dataset. (**B**) Further division of data using a clustering methodology. (**C**) Phylogenetic tree reconstruction and ancestral sequence reconstruction for the nodes on the trees. (**D**) Trend construction and amino acid prediction. (**E**) Prediction bundling.

The dataset of sequences obtained in previous steps is divided to create multiple phylogenetic trees of 150 leaves each. Thus, the number of trees depends on the size of the dataset. Each tree is ensured to have the target protein sequence. The dataset of sequences is parsed by ensuring that the headers cannot contain most of the special characters that are problematic (such as ( ) : ; . , or numbers headers starting with numbers) for several of the utilised tools. Next, SigClust (Liu et al., 2008) clusters sequences based on similarity. The parameter c is set to 150 to obtain up to 150 sequence clusters. Based on the clusters, a dataset of sequence files is constructed with the following conditions: (i) every sequence from the target set has to appear in at least one of the sequence files, (ii) one sequence is randomly selected from each cluster to construct one sequence file, (iii) to maximise diversity, the algorithm remembers which sequences from each cluster it already used and does not include them in the multiple sequence files unless it runs out of options (the number of sequence files will therefore be equal to the size of the largest cluster), and (iv) each sequence file has to contain target sequence. Once the sequence files are constructed, ClustalOmega (Sievers & Higgins, 2014) is used to construct a multiple sequence alignment (MSA) separately for each sequence file.

The MSA is subjected to the standard FireProt^ASR^ workflow (Musil et al., 2020). RAxML (Stamatakis, 2014) is employed to construct a phylogenetic tree for each MSA of the sequence files using the maximum-likelihood algorithm. RAxML is run in its SSE3 version using 50 bootstraps and the best-fitting evolution matrix suggested by IQ-TREE, most commonly parametrized as PROTGAMMALG. The resulting phylogenetic trees are amended by removing the evolutionary distance above the tree root. The minimum ancestral deviation algorithm is utilised to root the phylogenetic trees. Rooted phylogenetic trees and corresponding MSAs are then subjected to the LAZARUS (Hanson-Smith et al., 2010) calculation to obtain posterior probabilities for each sequence file. LAZARUS is run with the “codeml” module, corresponding evolutionary matrix, and fixed branch lengths. Gap reconstruction is disabled in LAZARUS as this step is done by the “gapCorrection” script implemented in FireProt^ASR^. Once the posterior probabilities and the ancestral gaps are calculated, ancestral sequences for each node in the phylogenetic tree are identified as the most probable amino acids/gaps. From the phylogenetic tree, the main path from the root to the target sequence is identified for each phylogenetic tree representing individual sequence files.

The target and sequences from all ancestral nodes between the target and the roo tare parsed into a separate file and aligned using ClustalOmega. Finally, a Python script using the “numpy” and the “sklearn.linear_model” libraries (*Sklearn.linear_model.LinearRegression*, 2023) is used to calculate the successor as the next step on the regression curve as follows: (i) For each column of multiple-sequence alignment (‘Trajectory’), the matrix of amino acid physico-chemical features is obtained. The matrix has nine columns, one from each of the selected AAindices, (ii) the vector of changes in the physico-chemical features, weighted by the distance from the root node, is calculated for each column of those matrices, (iii) this vector is consequently utilised to train the linear regression model to predict the next amino acid in the trajectory, thus mimicking laboratory-based protein evolution, (iv) the distance (between values of the AAindex for subsequent amino acids in a path) of the next step is calculated as an average distance of the nodes in the main path in the phylogenetic tree, (v) once the regression for each physico-chemical feature is calculated separately, these values are then used to assign categories and bundle predictions (**Table 2**), and (vi) the procedure is repeated for each column of multiple-sequence alignment and each sequence file.

**Table 2.**
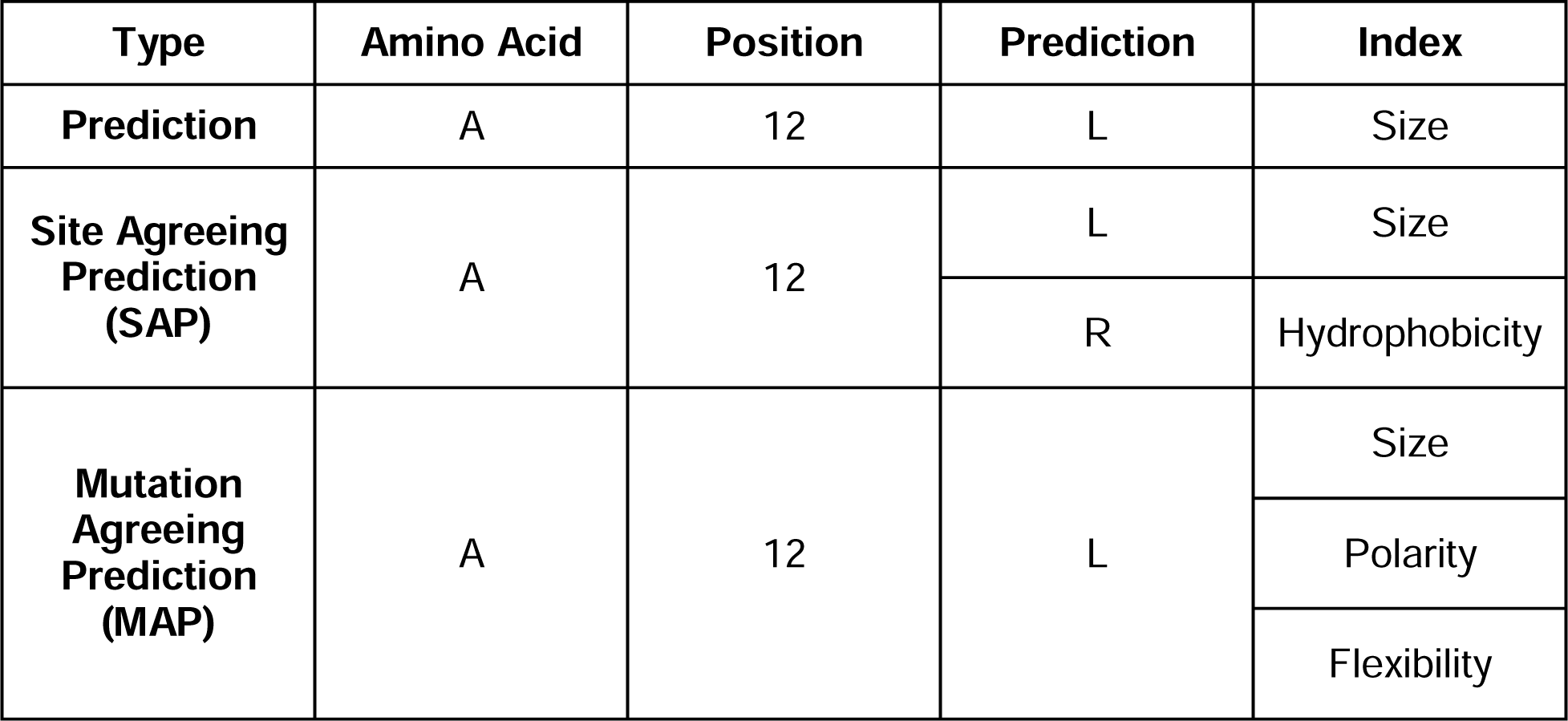
An example of the generalised prediction bundling scheme for three different levels of prediction: Prediction, Site Agreeing Prediction (SAP), and Mutation Agreeing Prediction (MAP)

Multiple measures are implemented to minimise the over-interpretation of a minimal linear regression-based strategy. The generated linear regression plot is divided by the number of transitions - one amino acid changing into another over a trajectory. Every time an amino acid does not change over a similar position over subsequent ancestors, it is treated as a grouping from the perspective of transitions and thus is not negatively scored, i.e., transitions between groupings are only counted.

The penultimate transition is important; if the change in transition does not agree with the overall trend, it is marked as a *‘break trend’*. The *sequentiality* of transitions of a trajectory is also scored out of 100. A score of 100 is achieved, for example, when in a positive trend, every subsequent transition has a higher value than the previous one. *Fluctuations* in a trajectory are scored using the amount of different amino acids divided by the number of amino acid groups. This demonstrates how much the trajectory ‘fluctuates’ or stays stable. No predictions are made for sites with less than three transitions.

These scores are used to rank successor amino acid predictions per site, per index, and per tree. The highest-scoring prediction should have high *sequentiality*, high *fluctuation,* and no *break trend* at the penultimate amino acid position. Each single amino acid prediction is averaged per site and tree. When multiple such predictions agree at the site but not at the mutation, they are termed Site Agreeing Predictions (SAPs). When multiple predictions from different AAindices agree on the mutation, the prediction is bundled into a single prediction. These predictions are termed Mutation Agreeing Predictions (MAPs) (**Table 2**).

### Validation Datasets

Various datasets were investigated as test cases for SSP. This includes homolog sets for levoglucosan kinase - UniProt ID B3VI55 (Klesmith et al., 2017), cold shock protein CspB - UniProt ID P32081 (Gribenko & Makhatadze, 2007), ADP-ribosylarginine hydrolase - Uniprot ID P54922 (Slodkowicz & Goldman, 2020), and aminoglycoside 3’-phosphotransferase - UniProt IS P00552 (Melnikov et al., 2014).

Individual datasets were compiled in different ways. The levoglucosan kinase set was found via the *in-house* SoluProtMut^DB^ database (Velecký et al., 2022) by searching for a protein with a large number of experimentally validated single-point mutations and their effects on the solubility of the protein. Similarly, the cold shock protein CspB dataset was found in the *in-house* FireProt^DB^ database (Stourac et al., 2020), by searching for a protein with a large number of experimentally validated single-point mutations and their effects on the thermostability of the protein. In cases where multiple values were available for a single mutation, the mean was taken. ADP-ribosylarginine hydrolase dataset was picked as it was one of the example cases for Slodkowicz & Goldman’s online tool (2020) for Structure Integrated with Positive Selection. ADP-ribosylarginine hydrolase was picked after a literature review, due to the sheer number of single-point mutations tested (fully site saturated) on the target protein by Melnikov et al., (2014). This naturally presented a perfect test case for SSP. Individual and detailed dataset handling steps are noted in <u>SI 2</u>.

## Results and Discussion

### Dataset Statistics

We tested the performance of SSP on the homolog sets for levoglucosan kinase (solubility), cold shock protein CspB (thermostability), ADP-ribosylarginine hydrolase (selectivity), and aminoglycoside 3’-phosphotransferase (activity). It is important to note that with the exception of Aminoglycoside 3’-phosphotransferase dataset, none of the other datasets used in the study have the values for the relevant effect for every possible point mutation that SSP predicts. Thus it is not possible to validate all predictions made by SSP. The results section only shows validation based on all mutational data points that SSP predicted and for which experimental labels were available. Figure 3 summarises the total single-point mutational space, the available experimental values, the number of predictions, and the overlaps between the two.

**Figure 3.**
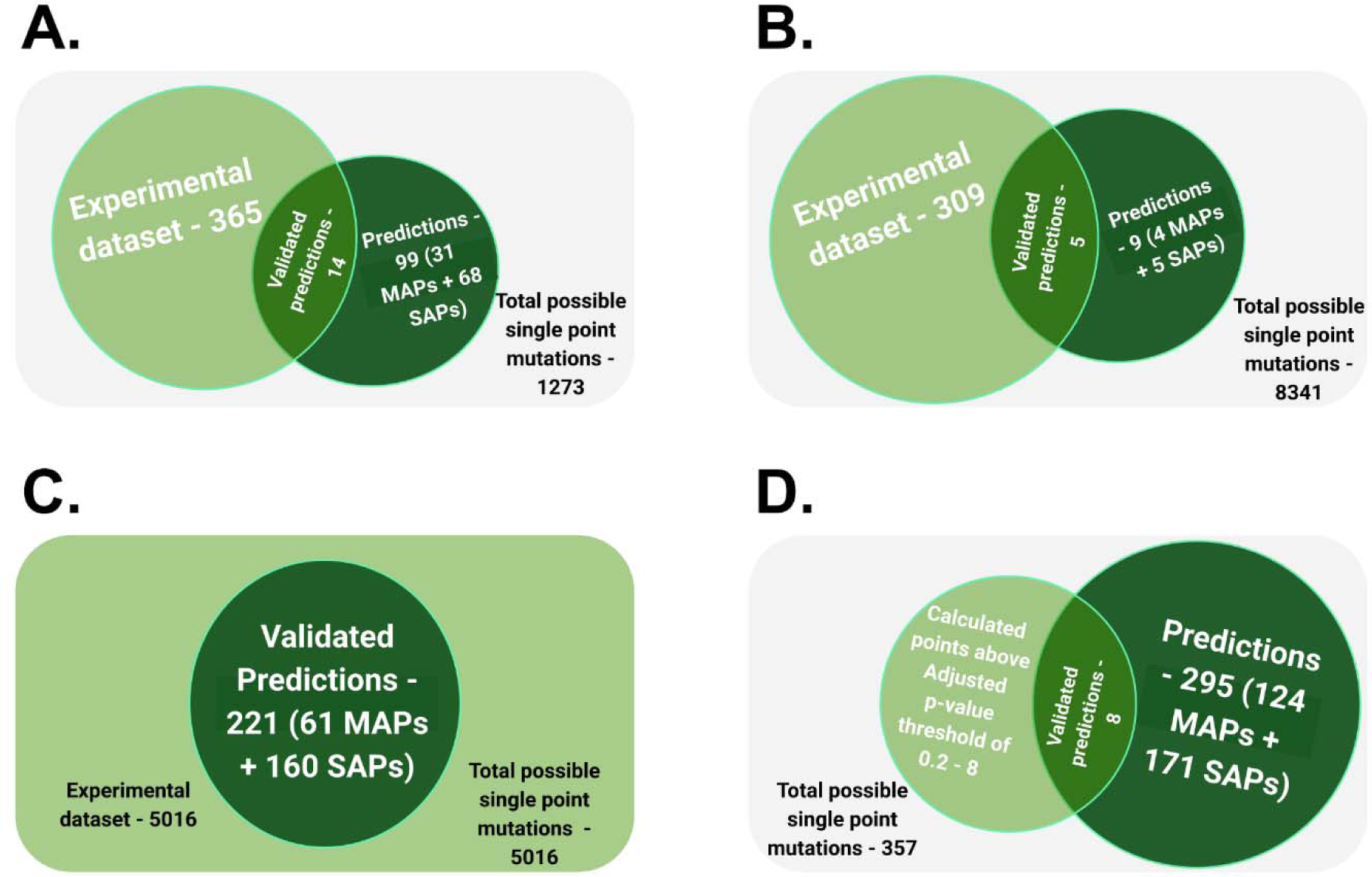
The visualisation of overlaps between the available experimental data and the predicted data. (**A**) Overlap metrics for Cold shock protein CspB set (FireProt^DB^ dataset - Stourac et al., 2020), (**B**) Overlap metrics for levoglucosan kinase set (Klesmith et al., 2017) (**C**) Overlap metrics for Aminoglycoside 3’-phosphotransferase set (Melnikov et al., 2014), and (**D**) Overlap metrics for ADP-ribosylarginine hydrolase set (Slodkowicz & Goldman, 2020). The experimental data are represented by a light green circle, while a dark green circle represents predicted data.

### Engineering Thermostability

SSP predictions for Cold shock protein CspB were compared to experimental data points with known effects of the mutation on protein thermostability from a collated dataset stored in the database FireProt^DB^ (Stourac et al., 2020). In cases where values from multiple datasets were available, the mean values were noted. E3Q was the only MAP that was supported by more than three indices. E3K was supported by 2 indices, and all others were SAPs. The results are provided in **Table 3**.

**Table 3.** Effects of mutations generated by SSP on the thermostability of Cold shock protein validated against the collated FireProt^DB^ dataset (Stourac et al., 2020)

| Mutation by SSP | Mean $\Delta\Delta G$ (kcal/mol) | $\Delta T_m$ (°C)* | Prediction agreement type |
| --- | --- | --- | --- |
| L2R | +0.4 | N/A | Site agreeing |
| E3K | -2.5 | +16.6 | Mutation agreeing |
| E3Q | -1.1 | N/A | Mutation agreeing |
| E3R | -1.7 | +16.0 | Site agreeing |
| E3V | -1.8 | N/A | Site agreeing |
| D24N | +0.7 | N/A | Site agreeing |
| A46E | +0.1 | -5.0 | Site agreeing |
| A46K | -1.4 | +8.4 | Site agreeing |
| A46L | -0.8 | N/A | Site agreeing |
| E50K | +0.3 | N/A | Site agreeing |
| N55D | -0.5 | N/A | Site agreeing |
| N55K | 0.0 | +0.8 | Mutation agreeing |
| N55S | +0.2 | N/A | Site agreeing |
| E66K | -2.2 | +12.9 | Site agreeing |

\*N/A - data not available.

There were 365 total mutations in the FireProt^DB^ dataset (Stourac et al., 2020), of which 18% were enhancing mutations in terms of thermostability (ΔΔG lower than −1 kcal/mol), 55% were neutral (ΔΔG from −1 kcal/mol to 1 kcal/mol), the remaining 27% were destabilising (ΔΔG greater than 1 kcal/mol). Any value between these ranges was considered neutral. The thresholds for stabilising, neutral and destabilising values were taken from the FireProt^DB^.

From the 14 mutations predicted by SSP, six were stabilising. The other eight mutations had ΔΔG values between −1 kcal/mol and 1 kcal/mol and can thus be classified as neutral. Five out of 14 mutations also increased the melting temperature - Tm, of the protein, and only one was destabilising (**Table 3**).

### Engineering Solubility

SSP predictions for levoglucosan kinase were compared to experimental data from Klesmith et al. (2017) available in the SoluProtMutDB (Velecký et al.). This comparison assessed how well the SSP predictions matched the known effects of mutations on protein solubility.The mutations I3L and I3F (supported by two different indices) had a neutral effect on solubility. Both mutations predicted by SSP, D9G and K38Q, are known to have a slightly enhancing effect on solubility. Only V200A showed a slightly negative effect on solubility in *E. coli* (**Table 4**). This suggests that the expressed mutants produced via SSP do not compromise their solubility.

**Table 4.**
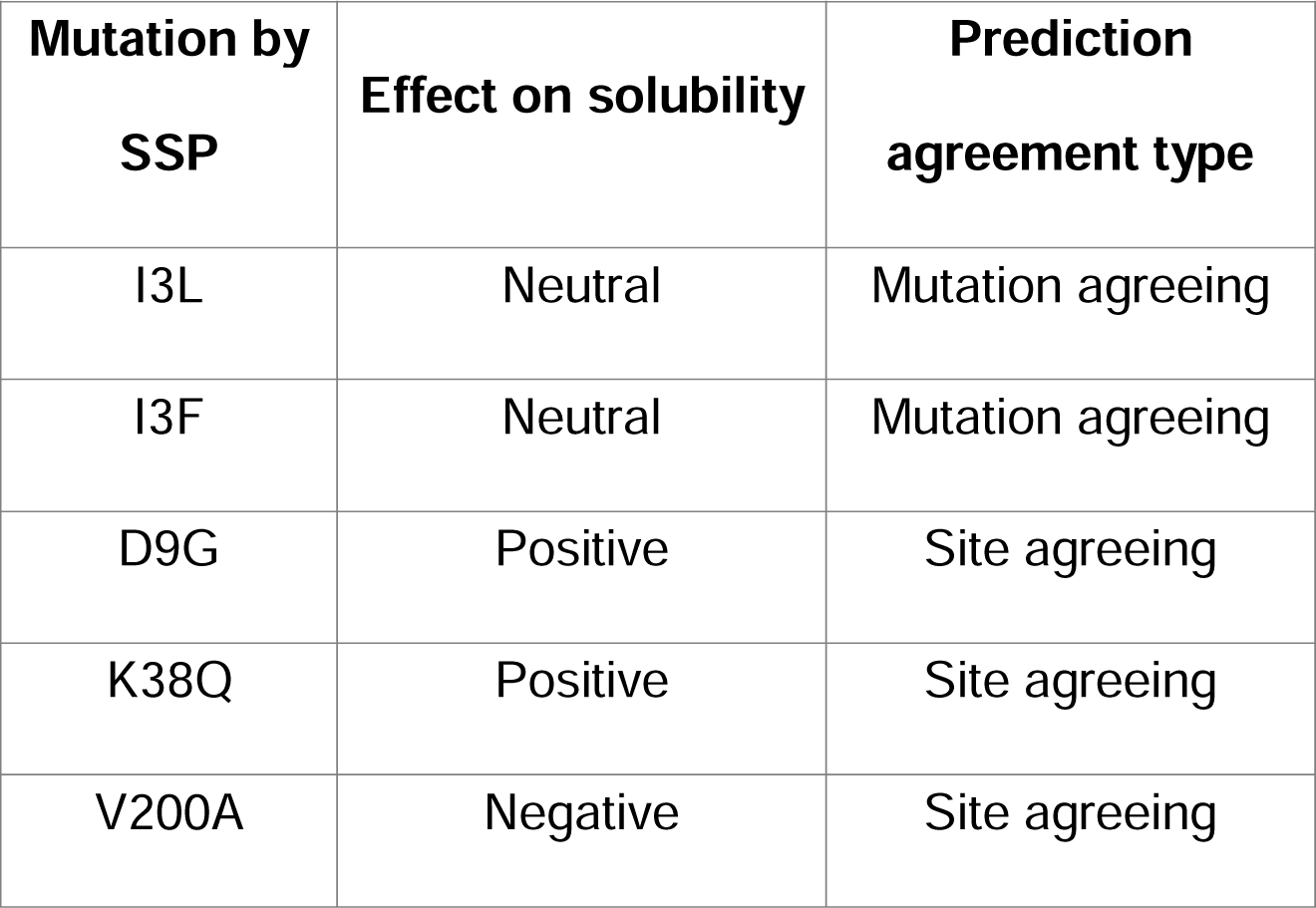
Effects of mutations generated by SSP on the solubility of levoglucosan kinase (Klesmith et al., 2017)

| **Mutation by SSP** | **Effect on solubility** | **Prediction agreement type** |
| --- | --- | --- |
| I3L | Neutral | Mutation agreeing |
| I3F | Neutral | Mutation agreeing |
| D9G | Positive | Site agreeing |
| K38Q | Positive | Site agreeing |
| V200A | Negative | Site agreeing |

### Engineering Activity

Aminoglycoside 3’-phosphotransferase is a protein that confers resistance to aminoglycosides, such as kanamycin, neomycin, paromomycin, ribostamycin, butirosin, and gentamicin B. Melnikov et al. (2014) conducted site saturation mutagenesis on this protein, transformed variants into cells and exposed them to six different antibiotics at up to four different concentrations. The amino acid enrichment (the number of identified variants with the particular mutation) was then noted in each case. A value of ∼1 applies to wild types, while a higher value means more resistance and hence more significant enrichment of that mutant, and *vice versa* for a value below 1 (Figure 4).

**Figure 4.**
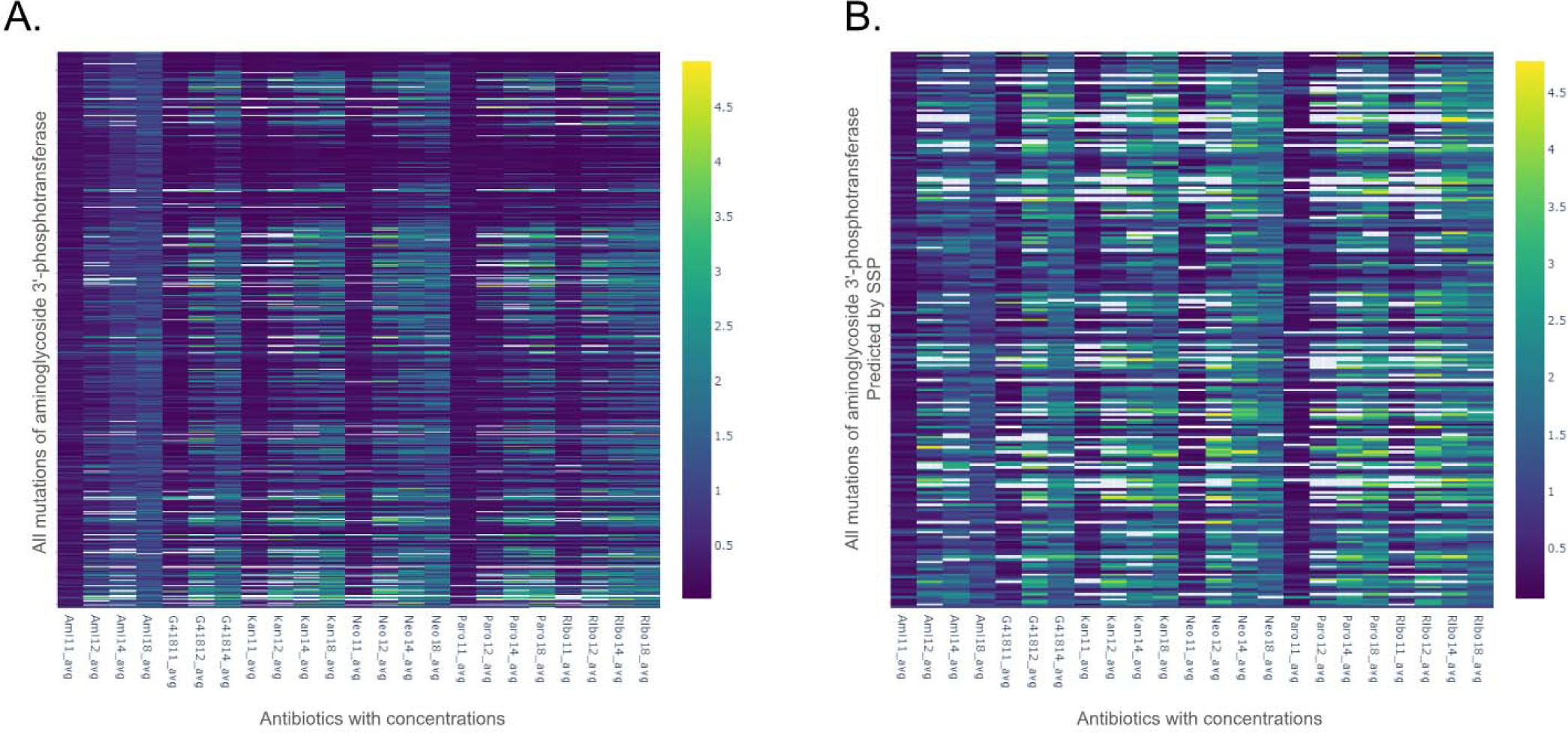
Heatmap visualisations comparing the enrichment values for mutations of aminoglycoside 3’-phosphotransferase. (**A**) A heatmap representing the entire mutational space of aminoglycoside 3’-phosphotransferase. (**B**) A heatmap representing only the mutations of aminoglycoside 3’-phosphotransferase that were predicted by the SSP. The X-axis represents the antibiotics and their tested concentrations, while the Y- axis represents the relevant mutations of aminoglycoside 3’-phosphotransferase. Details of antibiotic concentrations and individual enrichment values can be found in <u>SI 1</u>. Each rectangle on the plot indicates the enrichment value for a mutation when exposed to the effects of the specific antibiotic concentration. The Viridis colour map is used to maintain perceptual uniformity. A value of 1 (dark blue) represents no change in enrichment from the wild type, anything below 1 (purple) represents a negative effect on enrichment, while anything above 1 (light blue to yellow) represents a positive enriching effect of the mutation. This figure contrasts the effects of random mutations on the activity of aminoglycoside 3’-phosphotransferase, against the effect of SSP suggested mutations for the same protein. The perceptual increase in ‘brightness’ of Figure 4 B over Figure 4 A illustrates an increase in the positive impact of mutations on the activity of aminoglycoside 3’-phosphotransferase.

The average of all enrichment values across antibiotics and their concentrations in the complete dataset (AAC value) was 0.82. This means that a random, single-amino- acid variant is less likely to be resistant than the wild type, and, therefore, will have lower activity. The AAC value of 0.82 may be assumed as a proxy value for random mutations, while 1 is the default value for the wild type. Thus, random single-point mutations are likely to reduce the protein’s fitness. SSP generated 221 predictions, all with experimental validation points available from this large-scale site saturation mutagenesis study (Figure 4). For mutants generated by SSP, the AAC value is 1.36, showing a preferable selection of enriched (more active) variants, thus an increase in fitness if a mutation is selected from SSP’s output. Moreover, 61 of the 221 mutations were predicted at the MAP level, and their AAC value is 1.4. The remaining 160 predictions were made at the SAP level, and their AAC value is 1.36. As the AAC value for MAP level predictions is slightly higher than that for SAP level (1.4 *versus* 1.36), it hints at the possibility that MAPs may be slightly more reliable. This is summarised in Figure 3 C. The comparison of experimentally determined and predicted values are available in the supplementary table <u>SI 1</u>.

### Evolutionary Selection

Structure Integrated with Positive Selection (SIPS) is an online resource with positively selected sites mapped onto protein structures from an evolutionary perspective (Slodkowicz & Goldman, 2020). ADP-ribosylarginine hydrolase, which is one of the example cases of SIPS, has eight positively selected sites with an adjusted p-value threshold of 0.2 or higher. SSP predictions were made for ADP-ribosylarginine hydrolase to see how many of the predictions could be made for positively selected sites. Here, the emphasis was on sites and not the mutation itself, as SIPS only lists sites of evolutionary interest and not what they would mutate into. Out of the eight sites, seven were predicted by SSP, and six were MAPs, implying that SSP can selectively make predictions for sites with evolutionary significance.

**Table 5.** Cross-matching positively selected site data of ADP-ribosylarginine hydrolase from SIPS with SSP predictions (Slodkowicz & Goldman, 2020)

| **Sites selected**<br>**by SSP** | **Adjusted p-**<br>**value** | **Prediction agreement**<br>**type** |
| --- | --- | --- |
| K72 | 0.0848 | Mutation agreeing |
| P74 | 0.0473 | Mutation agreeing |
| T77 | 0.1731 | Mutation agreeing |
| Q78 | 0.1101 | Mutation agreeing |
| Q109 | 0.0796 | Mutation agreeing |
| H145 | 0.05 | Non agreeing |
| L189 | 0.0002 | Mutation agreeing |
| I355 | 0.0128 | *Not predicted* |

## Discussion and conclusions

SSP is a technique that allows the prediction of the evolution of amino acids in a protein sequence. It builds a statistical, ancestral sequence reconstruction-guided evolutionary history of a protein sequence (Spence et al., 2021), which is utilised to extrapolate the possible *future* substitution at a given position. SSP makes these predictions in the context of AAindex scoring (Kawashima, 2000) applied to the reconstructed evolutionary history for each position in a protein sequence. The AAindices used for SSP have been manually selected to reflect a variety of possibly relevant physiochemical descriptors. The selected set of AAindices can be easily adjusted based on the physico-chemical properties expected to be involved in shaping the evolution of a particular protein.

It should also be noted that while SSP utilises ASR, they are both *fundamentally different* techniques with distinct goals. ASR purports to ‘look back’ into the evolutionary history of a protein sequence, while SSP is designed to extrapolate into the potential future of a protein sequence. ASR is generally used for evolutionary analysis (Cai et al., 2004) and protein engineering (Spence et al., 2021). While ancestral proteins are more robust and with unique substrate specificities (Babkova et al., 2017; Hanson-Smith et al., 2010; Risso, V. A., & Sanchez-Ruiz, 2017), the engineering scope of ASR is generally along the lines of improving the thermostability of a protein and its expression yield. This is because ancestral proteins, when resurrected, tend to be more robust (Thomson et al., 2022). SSP can map out potential future evolutionary trajectories of a protein, and it can also be used to engineer proteins. However, it is not limited to improving thermostability and can amplify multiple other key properties of a protein as shown in this paper.

Proseeker is another tool that simulates natural selection and thus mimics evolution *in silico*. It uses physicochemical characteristics and structural information to pick sequences that have higher probabilities of evolving a desired function (Raven et al., 2022). However, the technique was designed specifically for binding proteins and lacks general applicability. Instead of complete protein sequences, it uses small peptide sequences (13 amino acids), and it also does not filter or select specific AAindices, rather it uses all available AAindices (Kawashima, 2000). The selection of relevant indices and then estimating their utility for any tool in this domain is crucial as many indices are redundant, e.g, nine indices for the hydrophobicity: ARGP820101, GOLD730101, JOND750101, PRAM900101, ZIMJ680101, PONP930101, WOLR790101, ENGD860101, and FASG890101 (Kawashima, 2000). This can lead to index weighting issues, where a certain physiochemical descriptor may have an exaggerated effect on the outcome. Furthermore, many indices are context-specific, such as hydrophobicity coefficients in specific solutions - from WILM950101 to WILM950104, and weights for alpha-helix at specific window positions - from QIAN880101 to QIAN880139 (Kawashima, 2000). Thus a careful selection of indices is a necessary step; SSP used manually curated non-correlated indices (**Table 1** and Figure 1). While the direct comparison between SSP and Proseeker could have been useful, it is hard to achieve as Proseeker works with shortened peptides (13 AA long) instead of the whole protein sequence. Moreover, it specifically requires binding affinity data to score every iteration of *in silico* evolution, thus making the technique specific to nucleic acid binding peptides.

SSP was validated using the datasets from different sources to test for the performance of various properties. In the case of thermostability, SSP made 14 predictions for the cold shock protein CspB, eight of which had a stabilising effect on the protein (ΔΔG < 0), while the remaining six were neutral with ΔΔG values between 0 kcal/mol and 1 kcal/mol. Five of the predicted mutations also had positive experimentally determined changes in melting temperatures LTm (°C), including the highest increase in melting temperature of +16.6 °C, and only a single mutation with a negative LTm (°C) value of −5°C.

SSP was also used to make predictions for aminoglycoside 3’-phosphotransferase (Melnikov et al., 2014). Aminoglycoside 3’-phosphotransferase is an enzyme that confers resistance to aminoglycosides with antibiotic properties. Thus an enhancement of enzyme’s activity can increase the antibiotic resistance of a bacteria that codes for it. SSP made 221 predictions for Aminoglycoside 3’-phosphotransferase with an AAC value of 1.4 at the MAP level, and 1.36 at the SAP level (1 being the value for the wild type, and 0.82 being the average value for random mutagenesis), thus demonstrating predictive prowess in the context of enhancing enzymatic activity, being significantly better than random mutation, while conferring an improvement over the wild type itself.

Validation of mutations predicted from the solubility dataset showed a higher likelihood of a positive or neutral effect on the solubility of the protein, despite the sparseness of the dataset. Furthermore, evolutionary selectivity data for ADP-ribosylarginine hydrolase (Slodkowicz & Goldman, 2020) taken from SIPS and SSP made predictions for 7 of 8 evolutionary selected sites with an adjusted p-value upper threshold of 0.2. This result suggests that SSP is selective in making predictions for sites that tend to evolve under positive selection, thus making a strong case for SSP’s selectivity.

Analysis and validation of methods like SSP are challenging as gleaning useful insights from various datasets with differing standards can be complex due to limited overlaps between the experimental and the predicted set (Figure 3). However, this study shows that the approach enhances specialised proteins by predicting mutations that improve desired properties, such as thermostability, activity, and solubility. Crucially, it also shows that SSP does not make predictions for sites randomly, but picks sites that are known to evolve under positive selection. In general, SSP method will work better with the proteins under stronger selection evolutionary pressure. Further validation of the predictor with diverse protein structures is desirable to define applicability for protein engineering applications.

As the service to the community, we are now integrating SSPas a new module into the easy-to-use web server FireProtASR (https://loschmidt.chemi.muni.cz/fireprotasr/), which will make predictions accessible to non-experts, jointly with related strategies Ancestral Sequence Reconstruction (ASR) and generation using Variational Autoencoder (VAE).

## Supplementary Materials

<u>SI 1</u> - Comparison of experimentally determined and predicted values for Aminoglycoside 3’-phosphotransferase, as well as the complete enrichment data

<u>SI 2</u> - A text README file on how to locally run the Successor Sequence Predictor

<u>SI 3</u> - Example case query sequence in FASTA format

<u>SI 4</u> - Example case homolog database in FASTA format

<u>SI 5</u> - Pre-run Ancestral Sequence Reconstruction output

## Data and Code Availability

We provide scripts as a command line application written in Python 3.8, which can be found on GitHub https://github.com/loschmidt/successor-sequence-predictor. We use the LinearRegression module from the scikit-learn library to predict the evolutionary trend in the phylogenetic tree.

## Acknowledgements

This work was supported by Operational Programme Research, Development and Education - “Project Internal Grant Agency of Masaryk University” (No. CZ.02.2.69/0.0/0.0/19_073/0016943) and Brno University of Technology [FIT-S-23- 8209]. Authors thank to the RECETOX Research Infrastructure (No. LM2023069) financed by the Ministry of Education, Youth and Sports and National Institute for Cancer Research (Programme EXCELES, ID Project No. LX22NPO5102) - Funded by the European Union - Next Generation EU. Computational resources were provided by the e-INFRA CZ and ELIXIR-CZ projects (ID: LM2018140 and LM2023055), supported by the Ministry of Education, Youth and Sports of the Czech Republic. This project was also supported by the European Union’s Horizon 2020 Research and Innovation Programme under grant agreement No. 857560 and by the Czech Ministry of Education, Youth and Sports, and the Operational Programme Research, Development and Education (the CETOCOEN EXCELLENCE project No. CZ.02.1.01/0.0/0.0/17_043/0009632). Pavel Kohout is holder of the Brno Ph.D. Talent scholarship funded by the Brno City Municipality and the JCMM.

