## Supplementary material for "Anticipating Protein Evolution with Successor Sequence Predictor": SI 2

### SI2: How to locally run the Successor Sequence Predictor

This document contains the basic information required to locally run the Successor Sequence Predictor pipeline with the example case of levoglucosan kinase. The pipeline consists of both pre-existing tools as well as custom python scripts.

1. Upon identification of a target protein, a curated homolog dataset is established through BLAST, maintaining query identity within the 50-90% range and enforcing a length filter between 80%-120% of the target's length. For our purposes we will use the levoglucosan kinase sequence (**SI 3**), and its homolog dataset (**SI 4).**

**Note -** Steps 2 through 5 can be done manually by using the tools listed below, however as some of these tools require time to set up and some prior knowledge, **SI 5** has been provided which already has the required processed files. These files can be directly used with the python script used in step 6, available at *https://github.com/loschmidt/successor-sequence-predictor*

2. Clustering is performed on the entire dataset via SigClust.
3. Individual files of 150 sequences each along with mandatory inclusion of the target sequence are created. The sequences are picked randomly from different clusters to ensure diversity inside each set. Attempt is also made to ensure that each sequence from the original set shows up in at least one of the trees, while also minimizing repetition of the same sequence in different trees. This may not always be possible depending on the nature of the dataset. To simply replicate this step, use the *create_sets.py* script from the aforementioned repository linked above and follow the instructions in the README file.
4. Multiple-sequence alignments for each of these files are generated using ClustalOmega.
5. This is followed by the application of the FireProt-ASR workflow, involving RAxML for phylogenetic tree construction and LAZARUS for posterior probability calculation, to generate ancestral sequences
6. Traces are created for each amino acid site, from the ancestral root node to the extant sequence, for every tree and every index. This is done using our Python script (available at *https://github.com/loschmidt/successor-sequence-predictor*), which performs linear regression to predict successor amino acids based on physico-chemical features. Scores, such as sequentiality, fluctuation, and trend consistency, as discussed in the paper, are also generated using the same script. The script then uses these scores to bundle predictions at various agreement levels. The generated successor mutations may need to be renumbered to align with an extant reference sequence, which will have to be done manually.
